# Polyphasic circadian neural circuits drive differential activities in multiple downstream rhythmic centers

**DOI:** 10.1101/2022.10.11.511837

**Authors:** Xitong Liang, Timothy E. Holy, Paul H. Taghert

## Abstract

Circadian clocks align various behaviors such as locomotor activity, sleep/wake, feeding, and mating to times of day that are most adaptive. How rhythmic information in pacemaker circuits is translated to neuronal outputs is not well understood. Here we used brain-wide, 24-hr *in vivo* calcium imaging in the *Drosophila* brain and searched for circadian rhythmic activity among identified clusters of dopaminergic (DA) and peptidergic neuroendocrine (PNE) neurons. Such rhythms were widespread and imposed by the PERIOD-dependent clock activity within the ∼150 cell circadian pacemaker network. The rhythms displayed either a Morning, an Evening, or Mid-Day phase. Different sub-groups of circadian pacemakers imposed neural activity rhythms onto different downstream non-clock neurons. Outputs from the canonical M and E pacemakers converged to regulate DA-PPM3 and DA-PAL neurons. E pacemakers regulate the Evening-active DA-PPL1 neurons. In addition to these canonical M and E oscillators, we present evidence for a third dedicated phase occurring at Mid-Day (MD): the l-LNv pacemakers present the MD activity peak and they regulate the MD-active DA-PPM1/2 neurons and three distinct PNE cell types. Thus, the *Drosophila* circadian pacemaker network is a polyphasic rhythm generator. It presents dedicated M, E and MD phases that are functionally transduced as neuronal outputs to organize diverse daily activity patterns in downstream circuits.

## Introduction

Animals display daily rhythms in a variety of physiological processes and behaviors, such as locomotor activity, sleep/wake, feeding, and mating behaviors (1; 2). Many such rhythms are controlled by circadian timing mechanisms and they exhibit a variety of phases throughout the solar day. Furthermore, the daily spectrum of circadian phases is itself regulated by daily changes in the presentation of environmental zeitgebers, especially light and temperature.. Under laboratory conditions, the locomotor activity of the fruit fly *Drosophila* peaks twice a day, in the morning and in the evening. During the morning peak, the fly shows a daily peak of feeding behavior (3) and mating behavior (4; 5). Following each of these two activity peaks, the fly exhibits two separate sleep bouts: one around mid-day and the other generally throughout the night. These behavioral rhythms are driven by synchronous clock gene oscillations (molecular clocks) in ∼150 circadian pacemaker neurons (6). How a small population of circadian neurons, sharing a mono-phase molecular clock, regulates all the different phases of behavioral rhythms required for fitness of a species remains poorly understood.

Previously we reported that molecular clocks generate circadian neural activity rhythms with diverse phases among five major circadian neuron groups (7). Each group peaks at a specific time of day. Three laterally-localized circadian neuron groups: s-LNv, l-LNv, and LNd display spontaneous activity peaks in the morning (M), at mid-day (MD), and in the evening (E), respectively. Two dorsally-localized circadian neuron groups the DN3 and DN1 cells are sequentially active during the nighttime (N1) and (N2). Genetic analyses revealed that the molecular clock dictates a default morning phase onto the pacemakers, but the polyphasic activity pattern ensues due to the delaying activities of light and neuropeptide modulation within the circadian neuron circuit (8). This reproducible series of phasic activity periods displayed across the circadian pacemaker network may therefore represent dedicated phasic timepoints by which different downstream output circuits could achieve temporal order. This general problem - how output circuits are regulated by circadian pacemaker circuits - is a fundamental problem in the field of biological rhythms.

We previously showed that the morning and evening phases (defined by activity peaks in the s-LNv and LNd/5^th^ s-LNv neurons respectively), drive in biphasic activity patterns in the Ring Neurons of the Ellipsoid Body (RN-EB) and in a subset of the PPM3 dopaminergic neurons (9). Both the RN-EB and PPM3 neurons are downstream neural circuits responding to clock signals to promote locomotor activity. Thus, the M and E phases of activity in the pacemaker circuit underlie authentic circadian phasic information that shapes premotor output to drive daily rhythmic locomotion.

Here we ask whether other phasic activity periods presented by the pacemaker circuit – those of the mid-day (MD - l-LNv group), the early night (N1 - DN3 group) and the late night (N2 - DN1 group) likewise direct daily rhythmic activity in downstream responsive non-pacemaker circuits. The problem could be approached by testing specific populations known to regulate different physiological and behavioral daily rhythms, as candidate responders of MD, N1 or N2 phasic information. For instance, a recent study by (10) revealed a candidate neural output circuit that mediates the N1 and N2 phasic information to promote nighttime sleep. However, more such candidate neural output circuits remain to be characterized (11; 12). As an alternative approach, we measured spontaneous daily activity patterns *in vivo* across two populations of chemically-defined neurons that are known to influence physiology and behavior – dopaminergic (DA) neurons and peptidergic neuroendocrine (PNE) neurons. We report that both populations exhibit daily periods of spontaneous activity whose phases align with either the M, the E or MD phases of circadian pacemaker neurons. We focus on MD phase regulation and present evidence from both experimental manipulations and from normal developmental progression to support the hypothesis that the MD activity phase is dictated by l-LNv pacemaker group activity. Together these findings extend the hypothesis that polyphasic timing information from the *Drosophila* pacemaker circuit has broad functional significance. It spreads widely and independently through parallel downstream pathways to generate phase-diverse patterns of physiological and behavioral rhythms.

## Results

### Daily neural activity rhythms of dopaminergic neurons

Several aspects of fly physiology and behavior, such as locomotor activity, sleep/wake, feeding, and mating behaviors are regulated by neuromodulatory systems in the fly’s brain. Prominent among these systems is the diverse collection of dopaminergic (DA) neurons (13;14). Therefore, we extended our previous measurements of spontaneous activity in the PPM3 DA neurons (9) and asked whether the neural activity of other DA neurons might also display temporal bias in the first day of constant darkness, following 12:12 LD entrainment. Using *TH (tyrosine hydroxylase)-GAL4*, we imaged five spatially-distinct clusters of DA neurons in the fly’s dorsal protocerebrum: PAM, PAL, PPL1, PPM1/2, and PPM3 (15). With the exception of the PAM, each cluster displayed a unique circadian-rhythmic Ca^2+^ activity pattern (Figure 1AB). Three clusters showed prominent single Ca^2+^ activity peaks, but at different times of day: PAL neurons peaked around dawn, followed by the PPM1/2 cluster, which peaked around mid-day, and later, the PPL1 peaked around dusk. As we previously reported (9), PPM3 exhibits a bimodal activity pattern, with a peak at dawn and a second at dusk. These distinct and stereotyped daily activity patterns in DA neuron clusters were confirmed by using two independent genetic drivers: *TH-C-GAL4* and *TH-D-GAL4* (Figure S1AB), which separately label largely non-overlapping DA neuron clusters. Together they recapitulate most of the *TH- GAL4* expression pattern (17). We also measured DA Ca^2+^ activities at high frequency (1 Hz) for short periods *in vivo* in brains exposed acutely at different times of day. We found that DA neurons, while at the peak of their daily slow Ca^2+^ fluctuations, also displayed a more dynamic fast Ca^2+^ activity than those at the trough time (Figure S1D-K). These observations were motived by high-frequency sampling of calcium fluctuations in pacemaker neurons (18). That study revealed that each of the five pacemaker groups exhibits co-phasic peaks of slow and fast Ca^2+^ oscillations that are mechanistically-distinct. Importantly, the spontaneous daily neural activity patterns of DA neurons were completely arrhythmic in circadian-defective *per^0^* mutant flies (Figure 1C and Figure S1C). Lastly, we noted clear alterations in spontaneous DA activity patterns in the absence of PDF signaling, which also speaks to the importance of the circadian pacemaker network (Figure 1D). In a severe *pdfr* mutant background, the PPM1/2 and PPL1 activities were undisturbed. However, the morning peak of the PPM3 cluster was lost and that of the PAL cluster was displaced to the evening phase. In summary, many neurons of the *Drosophila* dopaminergic system exhibit clock-dependent and cluster-specific spontaneous circadian neural activity patterns. Notably, the *TH*-gal4+ DA neurons do not express oscillatory clock proteins (19): together the findings suggest that diverse daily rhythms of DA neuronal activity are imposed by neuronal activity from the circadian pacemaker network.

**Figure 1.**
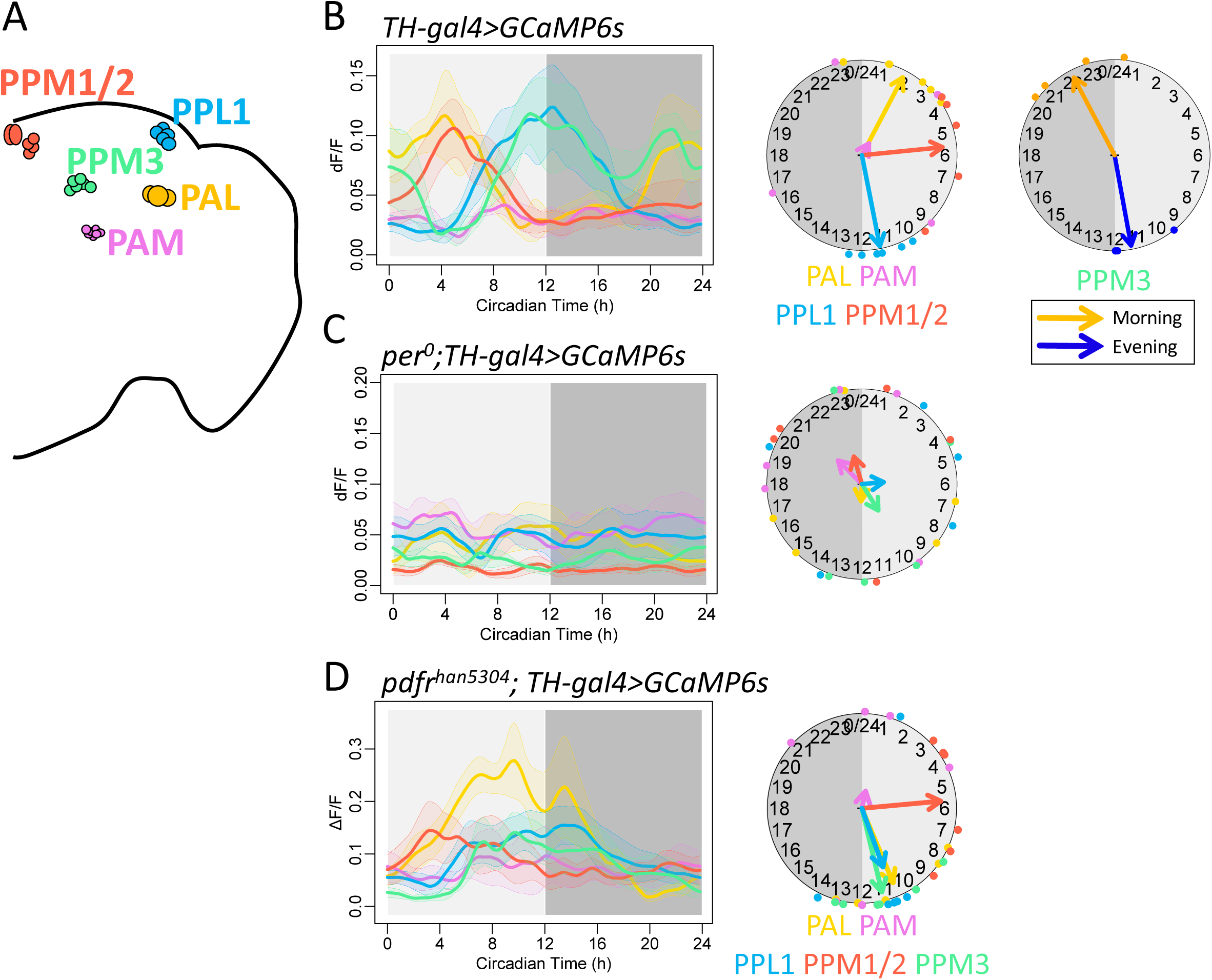
Diverse daily Ca^2+^ activity patterns of DA neuron clusters. (A) Map of the five DA neuron clusters accessible via in vivo imaging. (B) Left, daily Ca^2+^ activity patterns of DA neuron clusters under DD (n = 6 flies). Middle, Ca^2+^ phase distribution of DA neuron clusters showing single peaks (PAL, PPL1, and PPM1) or arrhythmic activity (PAM). Right, Ca^2+^ phase distribution of PPM3 for both morning peaks (orange) and evening peaks (blue). (C) Arrhythmic Ca^2+^ activity patterns of DA neuron clusters under DD in *per^01^* mutants (n = 5 flies). (D) Altered patterns of Ca^2+^ activity patterns of DA neuron clusters under DD in *pdf ^han5304^* mutants (n = 5 flies).

### Daily patterns of DA neurons in regulating sleep and mating behavior

We next compared the diversity of DA neuron daily activity patterns with the functional diversity of DA neuron clusters, as described by previous studies. DA plays a key role in sleep/wake regulation as a wake-promoting factor (20, 21, 22). To promote wake, a pair of PPL1 DA neurons (which represent a subset of the PPL1 group) project to the dorsal fan-shape body (dFSB) and suppress dFSB sleep-promoting neurons (17, 24, 25) Using highly specific, split- GAL4 drivers (13), we found that the specific PPL1-dFSB subset of PPL1 DA neurons showed a prominent evening Ca^2+^ activity peak (Figure 2AC), when the flies were most active under DD. Unlike the averaged Ca^2+^ peak from entire PPL1 cluster (Figure 1B), the Ca^2+^ activity of PPL1- dFSB subset was narrower and was centered on the phase of the evening locomotor activity peak (Figure S2). It dropped immediately at the beginning of subjective night, consistent with the onset of nighttime sleep. Thus, the specific Ca^2+^ activity pattern of wake-promoting PPL1-dFSB precisely matched the daily sleep/wake pattern around dusk. In addition to sleep/wake cycles, specific DA neurons also regulate courtship and mating behavior. The internal drive of male mating behavior is encoded by the activity level of a pair of *fruitless*-positive DA neurons from the PAL cluster (*Fru+* PAL, Figure 2B) (26). We found that restricting GCaMP6 expression to just these two identified *Fru+* PAL neurons revealed a spontaneous morning Ca^2+^ peak (Figure 2D), matching the daily behavioral mating activity pattern (4). Together, these observations suggest that two distinct subsets of DA groups are spontaneously active at different times of day. The different times align precisely with the phases displayed by the behaviors the neurons are known to help regulate. These baseline circadian activity patterns may therefore contribute to the mechanistic basis that differentially times modulation of diverse behavioral rhythms.

**Figure 2.**
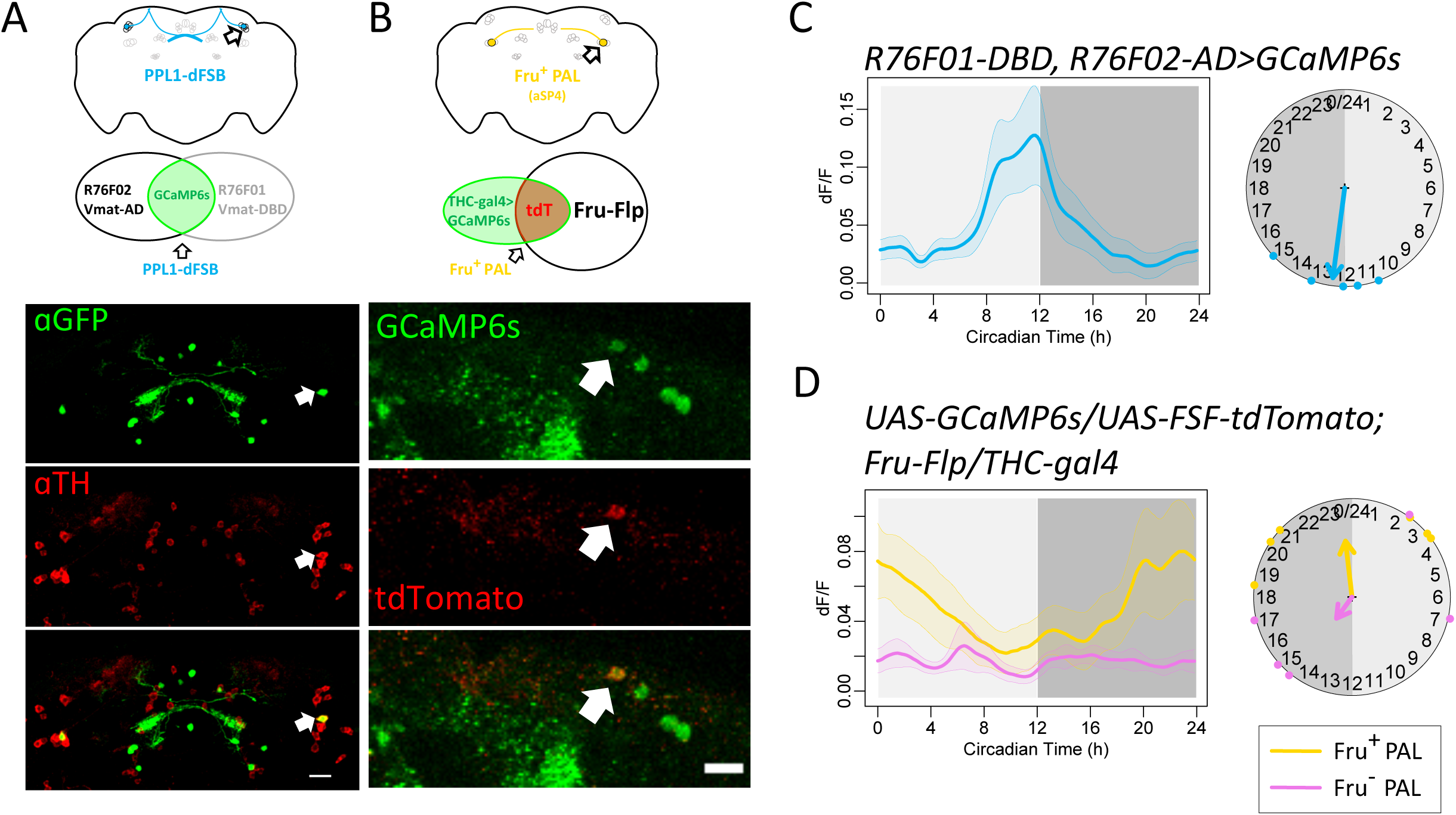
Daily patterns of DA neurons in regulating sleep and mating behavior. (**A**) Diagram and confocal images of a pair of PPL1 projecting to dorsal fan-shape body (dFB) neurons co-localized with immunostaining signal of tyrosine hydroxylase (TH). Scale bar, 20 μm. (Data re-plotted from 13) (**B**) Diagram and confocal images of a pair of fruitless positive PAL was labeled by tdTomato with intersection of TH-C-Gal4 and Fru-Flp. (**C**)Daily Ca^2+^ activity patterns of PPL1-dFB shown in (D) under DD (n = 5 flies). (**D**) Daily Ca^2+^ activity patterns of Fruitless-positive and -negative PAL shown in (E) under DD (n = 6 flies).

### Daily neural activity rhythms of peptidergic neuroendocrine neurons

Peptidergic neuroendocrine (PNE) cells release diverse peptide hormones and represent a second major neuromodulatory system (27). Many neuroendocrine cell bodies in insects localize in the *Pars Intercerebralis* (PI), a functional homolog of the mammalian hypothalamus (Figure 3AB). They have been implicated in regulating sleep (28) locomotion (30), and metabolism (31, 32). We identified these neuroendocrine cells by a peptidergic neuron marker DIMMED, which is a bHLH transcription factor associated with neuroendocrine cell differentiation (33). Using the *c929-GAL4 driver* that reports on the *dimm* promotor (33), we simultaneously imaged from all PI cells, from the pair of large laterally leukokinin neurons (35), and also from the DIMM-positive circadian neuron group, l-LNv (36). We found that on average, PI neurons displayed a daily Ca^2+^ activity peak around mid-day, with a phase very similar to that of the l-LNv (Figure 3C). PNE cells in the PI are heterogeneous: different neurons release different neuropeptides (33) which regulate different physiological and behavioral functions. We genetically dissected PI neurons for imaging based on the types of neuropeptides they released, including DH44 (dromyosuppressin (DMS), and SIFamide (SIFa; 32), and insulin-like peptides (dILPs). Most PI groups, when analyzed with these more specific Gal4 drivers, had activity peaks prominently at mid-day (Figure 3B, E-G), consistent with the averaged signal from the entire PI group (Figure 3C). We noted one exception: the insulin-producing cells (IPCs, labeled by *dILP2-GAL4*), which peaked in the morning (Figure 3D). This pattern is consistent with previous observations that IPC display a higher electrophysiological firing rate when recorded *in vitro* in the morning, than at other times of day (31). Other studies suggest they might be regulated by the outputs of M cells (37). IPC activity peaks in the morning, suggesting that the release of insulin-like peptides peaks in the morning, and could coincide with the peak phase of the fly’s daily feeding rhythm (3). Outside of the PI, a pair of neuroendocrine neurons releasing leucokinin (LK) has been showed to be involved in regulating locomotor activity rhythms (38) and feeding behavior (35, 39). We found the LK neurons showed a daily activity peak in the evening, different from the other PNE cells we studied, but consistent with the prediction that LK neurons are suppressed by the morning-active PDF neurons (38). Together, our results show that many PNE cells produce spontaneous circadian rhythms of neural activity and, like the DA neurons group described above, the rhythms exhibit diverse phases of peak activity (i.e., M, E, and MD). This is consistent with the hypothesis that the release of diverse peptide and protein hormones is under complex, polyphasic circadian regulation.

**Figure 3.**
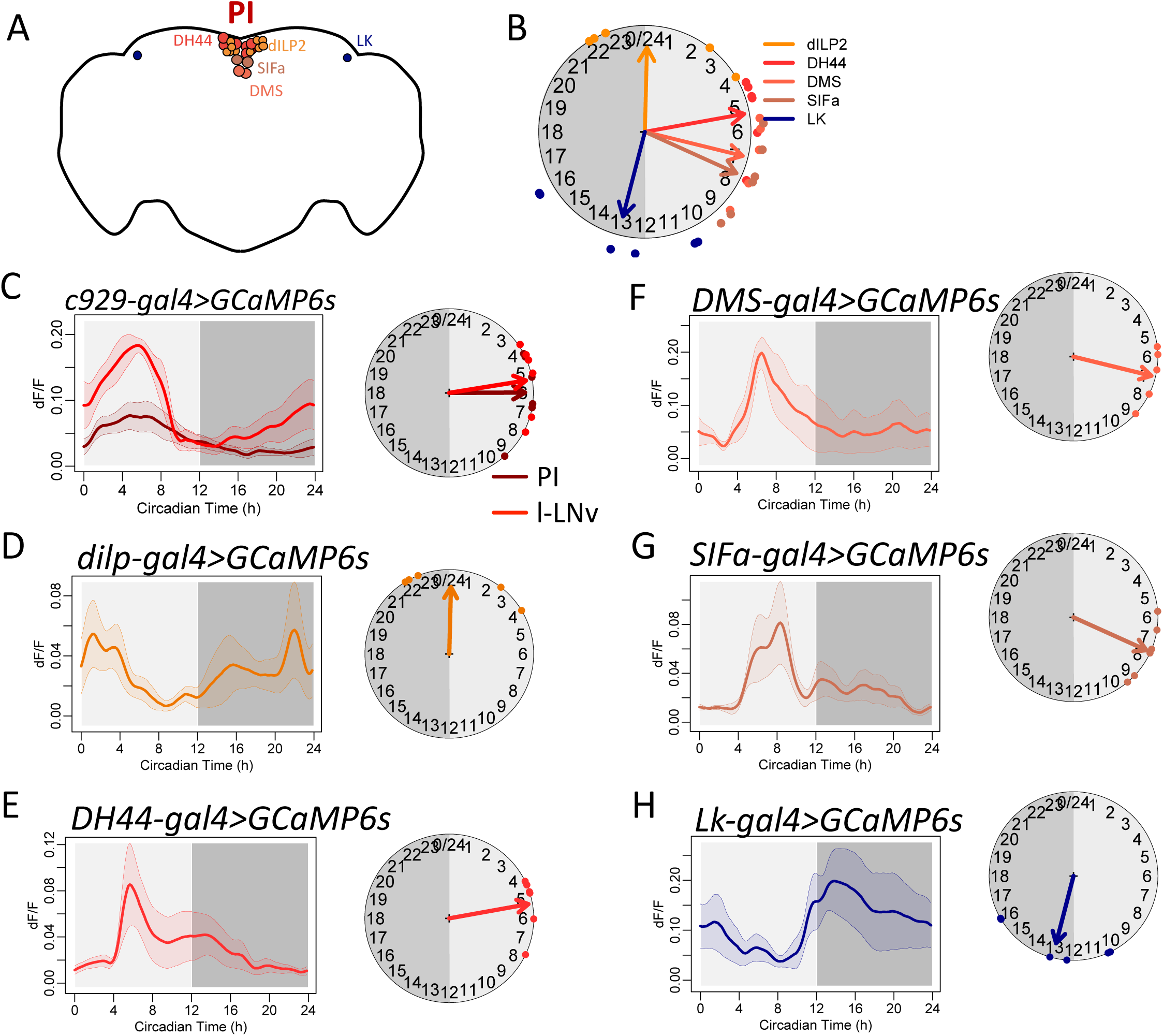
Daily neural activity patterns of neuroendocrine cells. (**A**) Diagram of peptidergic neurons in the brain. (**B**) Summary of phase distribution of different peptidergic neurons in (D-H). (**C**) Averaged Ca^2+^ activity rhythms of pars intercerebralis (PI) neuropeptidergic neurons and circadian neurons l-LNv labelled by *dimm(c929)-gal4* (n = 6 flies). (**D-G**) Daily neural activity patterns of four different of PI subgroups: insulin producing cells (labelled by *dilp2-gal4*), diuretic hormone 44 (DH44) neurons, dromyosuppressin (DMS) neurons, and SIFamide (SIFa) neurons (n = 5, 6, 5, and 6 flies). (**H**) Daily neural activity patterns of leucokinin (LK) neurons (n = 6 flies).

### Circadian neurons dictate phases of output circuits

The different output circuits we surveyed produced distinct phases of daily neural activity patterns. We developed several lines of evidence to test whether and how these patterns are temporally-organized by molecular clocks through the polyphasic outputs of the pacemaker network (7). We first asked whether different groups of circadian neurons regulate different downstream output circuits. For instance, do circadian neurons that peak in the morning generate corresponding morning output peaks? Likewise, do circadian neurons that peak at other phases of the 24-hr day produce corresponding co-phasic outputs? To do so, we selectively shift the activity phase of a subset, or even of single circadian neuron groups, by applying selective over-expression of Shaggy (SGG, *Drosophila* GSK3), a kinase for the clock protein TIM, that accelerates the molecular clocks when over-expressed (40). We measured the consequences by comparing the daily activity patterns of multiple output circuits. Driving SGG expression with *dvpdf-GAL4* (41) accelerated molecular clocks in M-pacemakers (s-LNv), MD-pacemakers (l-LNv), and E pacemakers (5^th^ s-LNv and LNds) (Figure 4A). In these flies, the Ca^2+^ peaks of M cells and E cells, as well as the morning and evening peaks of locomotor activity were advanced (Figure S3A and S3D). Corresponding to the behavioral phenotype, we found that the morning and evening peaks of PPM3 were significantly advanced (p = 0.029, 0.014; Watson-Williams test). Likewise, the daily activity peaks of PAL neurons in the morning and PPL1 neurons in the evening were also advanced (Figure 4A; PAL, p = 0.0097; PPL1, p = 0.028, Watson-Williams test). The phase of the Mid-Day PPM1 neurons was not affected.

**Figure 4.**
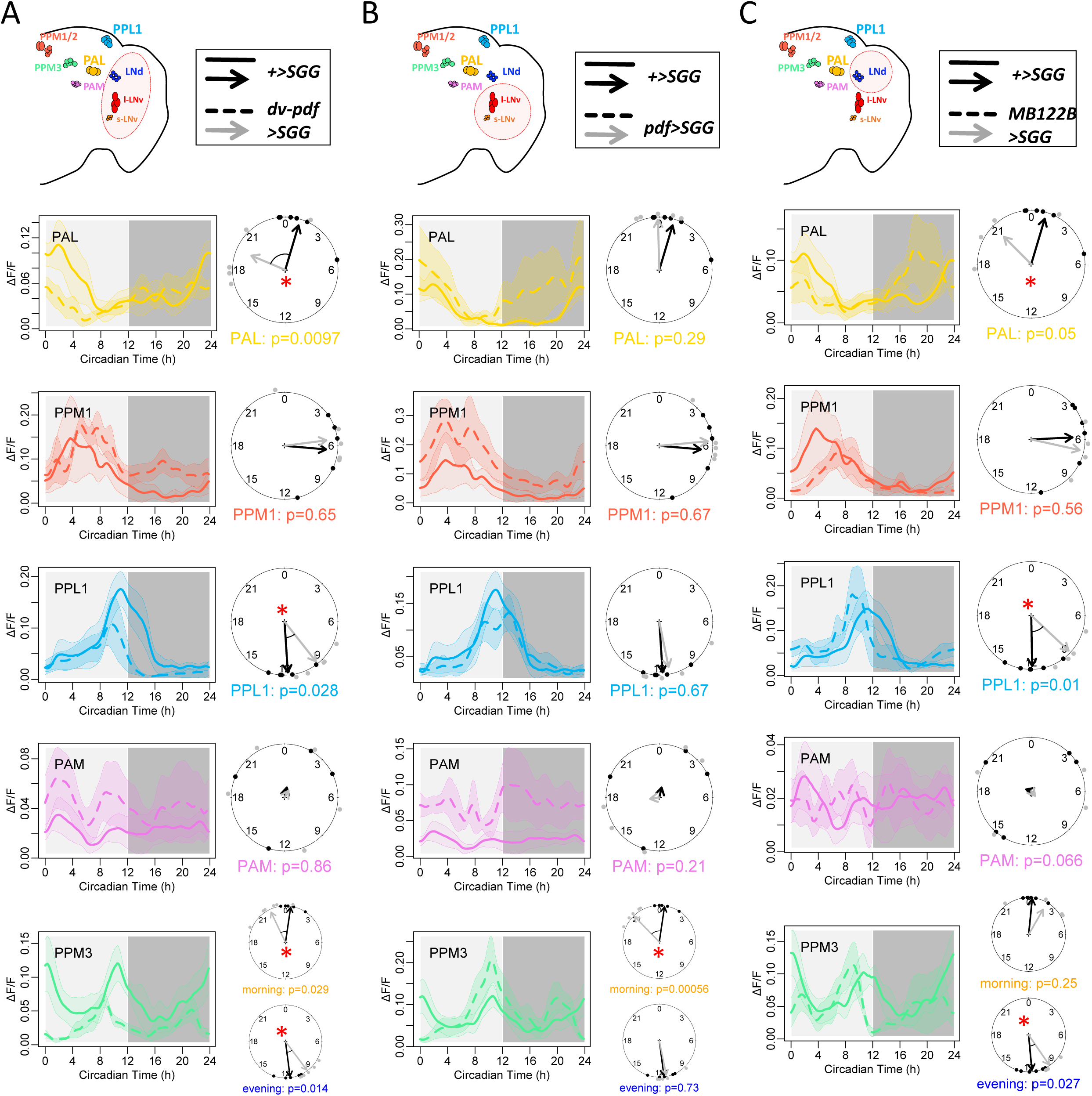
Daily activity phases of DA neurons PAL, PPL1, and PPM3 are dictated by M and E cells. (**A**) Daily Ca^2+^ activity patterns and phase comparison of DA neurons between WT flies under DD (solid lines, *+>SGG;TH-lexA>GCaMP6s,* n = 4 flies) and *dvpdf>SGG* flies under DD (dash lines, *dvpdf-GAL4>SGG;TH-lexA>GCaMP6s,* n = 6 flies). The daily peak phases of PAL and PPL1 and both morning and evening phases of PPM3 were advanced in *dvpdf>SGG* flies (* P < 0.05, Watson-Williams test). (**B**) Daily Ca^2+^ activity patterns and phase comparison of DA neurons between WT flies under DD (solid lines, n = 4 flies) and *pdf>SGG* flies under DD (dashed lines, *pdf-GAL4>SGG;TH- lexA>GCaMP6s,* n = 5 flies). Morning peak of PPM3 were significantly advanced in *pdf>SGG*. (bottom panel re-plotted from 9) (**C**) Daily Ca^2+^ activity patterns and phase comparison of DA neurons between WT flies under DD (solid lines, n = 4 flies) and *MB122B>SGG* flies under DD (dashed lines, *MB122B-splitGAL4s>SGG;TH-lexA>GCaMP6s,* n = 5 flies). The daily peak phases of PPL1, PAL, and the evening peak of PPM3 were advanced in *MB122B>SGG*.

We extended the analysis of changing clock phase in pacemaker subsets by using either of two more restrictive Gal4 lines. *pdf*-Gal4 restricted SGG over-expression to just the M and MD pacemakers: this caused a selectively advance of the M-cell (s-LNv) Ca^2+^ peak and the morning peak of locomotor activity (Figure 4B and Figure S3BE; 42). However, as found above with *dvpdf-GAL4,* the MD-cell (l-LNv) Ca^2+^ peak was not affected by over-expressing SGG. Among downstream circuits, we found that the morning phase of PPM3 neurons were selectively advanced (p = 0.00056; Watson-Williams test), while the PAL neuron morning phase was not. The second Gal4 line (the split-GAL4 driver *MB122B*) restricted SGG over-expression to the LNd and 5^th^ s-LNv E pacemakers. We observed a selectively advance in the E-cell (LNd) Ca^2+^ peak and in the evening peak of locomotor activity (Figure 4C and Figure S3CF; cf. 9). Likewise, the evening peaks of PPM3 neurons and the evening peaks of PPL1 neurons were both selectively advanced (PPM3, p = 0.027; PPL1, p = 0.01; Watson-Williams test). Unexpectedly, the morning peaks of PAL neurons were also advanced (p = 0.05; Watson-Williams test). Together, these results suggested that E cells may help signal proper activity phases for multiple downstream neuron groups, namely the PAL, PPL1, and the evening phase of the PPM3.

We were struck by the alignment of the l-LNv and PPM1/2 clusters, yet no SGG manipulation that we tested could alter the Ca*2+* activity phase of either of these MD-active cell groups. Therefore, to ask if the phase of l-LNv neuron activity can influence daily activity patterns of any downstream output neurons, we turned to l-LNv mis-expression of PDFR which delays its Ca^2+^ peak by as much as 6 hr (8). We used *pdf*-Gal4 to drive *pdfr* in the l- and s-LNv. This manipulation also slightly advanced the Ca^2+^ phase in s-LNv (Figure 5D), likely as a result of earlier termination of Ca^2+^ activation by autonomous PDFR (8). In downstream circuits, we found that the morning peak of PAL neurons was advanced (Figure 5E). PDFR over-expression also substantially delayed the Ca^2+^ peak in the l-LNv away from its MD phase by 5.7 hr (Figure 5D). Notably, the MD phase of the PPM1/2 DA neurons displayed a comparable multi-hr delay, such that they remained in synchrony with the l-LNv (Figure 5E). Thus PPM1/2 neurons are normally synchronous with the l-LNv at MD, and they remain aligned when the phase of the l-LNv is delayed as much as ∼ 6 hr by experimental manipulation.

**Figure 5.**
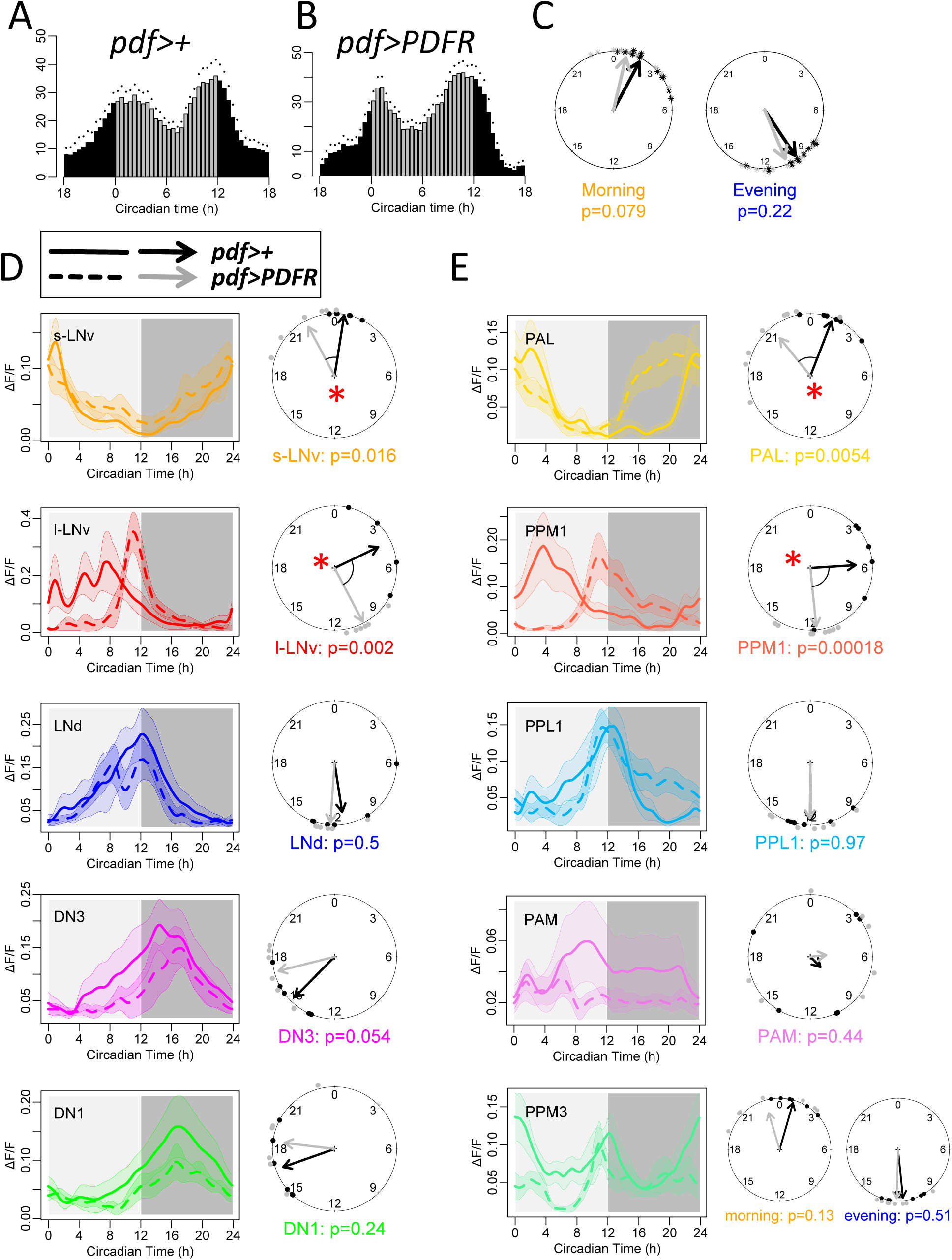
Daily activity phases of DA neuron PPM1 are dictated by circadian neurons l- LNv. (**A**) Average locomotor activity in DD1 of wild type flies (WT, n = 16). (**B**) Average locomotor activity in DD1 of flies expressing *PDFR* in PDF neurons using *pdf- GAL4* (n = 16 flies). (**C**) Phase comparisons of morning and evening activity between WT and *pdf>PDFR*. (**D**) Daily Ca^2+^ activity patterns and phase comparison of circadian neurons between WT flies (solid lines, *pdf-GAL4;cry-LexA>GCaMP6s*, n = 4 flies) and *pdf>PDFR* flies under DD (dashed lines, *pdf-GAL4>PDF;cry-LexA>GCaMP6s*, n = 5 flies). The daily peak phases of s-LNv were advanced while the daily peak phases of l-LNv were delayed in *pdf>PDFR* flies (* P < 0.05, Watson-Williams test). (**E**) Daily Ca^2+^ activity patterns and phase comparison of DA neurons between WT flies (solid lines, *pdf-GAL4;TH-LexA>GCaMP6s*,n = 4 flies) and *pdf>PDFR* flies under DD (dashed lines, *pdf-GAL4>PDF;TH-LexA>GCaMP6s*, n = 5 flies). The daily peak phases of PAL were advanced while the daily peak phases of PPM1 were delayed in *pdf>PDFR* flies (* P < 0.05, Watson-Williams test).

The l-LNv normally express little if any PDFR (19; 43), nor do they respond pharmacologically to PDF *in vivo* (11, 44). Likewise, loss of *pdfr* (*han* mutant flies) does not affect the normal l-LNv activity phase, whereas activity phases of the LNd and DN3 pacemakers are broadly phase advanced in the *han* background (7). Recently, Klose and Shaw (45) reported that on the first day of adult stage (the day of eclosion, termed Day 0), l-LNv express PDFR and respond pharmacologically to PDF. This transient period of PDFR expression declines by the third day to the low levels normally associated with the adult. We therefore measured the phases of Ca^2+^ activity of the entire pacemaker system and of selected downstream neurons on Day 0 to ask if the transient expression of PDFR (now due to a normal developmental progression, not an experimental consequence) had effects. We found that on Day 0, the s-LNv, LNd, DN1 and DN3 groups all displayed the same phases as found in mature adults, but that the phase of the l-LNv was delayed by 6.7 hr (Figure 6AB). Among the downstream DA and PNE neurons that we had measured in later adult stages, all maintained their normal phases, with the exceptions of those that normally peak at the MD phase. Ca^2+^ peak phases of PPM1/2 DA neurons and certain PI activity phases also displayed a substantial delay and so remained aligned with the l-LNv during this specific developmental stage (Figure 6C-F). Thus, during normal developmental progression at the earliest times in the adult stage, l-LNv transiently delay their activity peak as they transiently express PDFR. In conjunction, the activity peaks of downstream neurons that are normally synchronized with l-LNv at MD are also transiently delayed to the late day/early evening.

**Figure 6.**
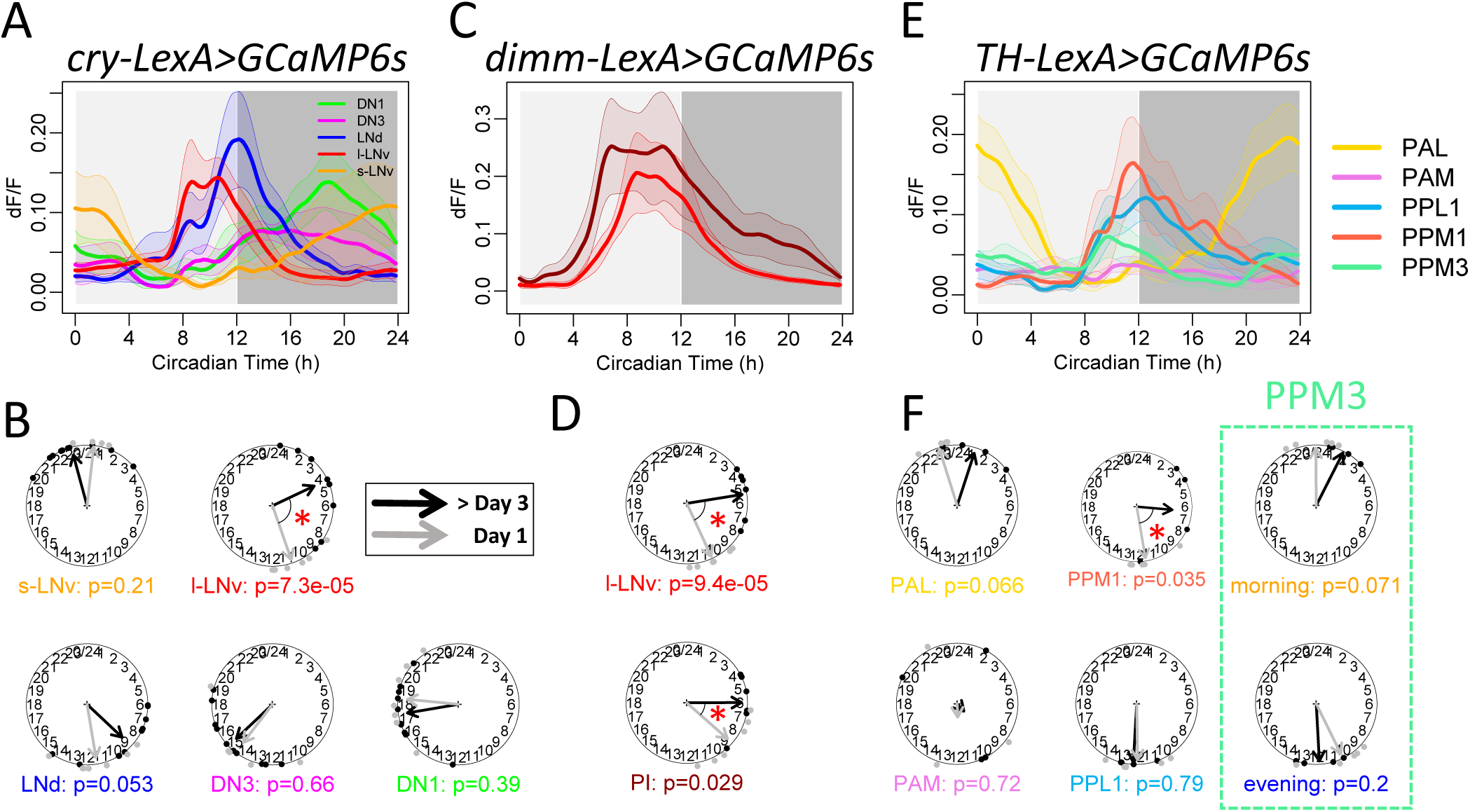
Daily activity patterns of circadian neurons and output circuits in the first day after eclosion. (**A**) Daily Ca^2+^ activity patterns of circadian neurons in flies within one day after eclosion (n = 7 flies). (**B**) Phase comparison between aged and newly eclosed flies. Note that l-LNv phases in newly eclosed flies were latter than that in aged flies. (*p < 0.05, Watson-Williams test). (**C**) Daily Ca^2+^ activity patterns of l-LNv and PI cells in newly eclosed flies (n = 6 flies). (**D**) l-LNv and PI cell phases in newly eclosed flies were latter than that in aged flies. (**E**) Daily Ca^2+^ activity patterns of dopaminergic neurons in newly eclosed flies (n = 5 flies). (**F**) PPM1 phases in newly eclosed flies were latter than that in aged flies.

Finally, we asked how pacemaker neurons communicate with non-clock downstream followers to shape their specific activity periods. We had already seen complex changes in spontaneous activity by the diverse DA neuronal groups lacking PDF signaling (*han* mutants, Figure 1D): the MD-phased (PPM1/2) and E-phased (PPL1) DA neurons sustained their normal times of activity. However, the PPMs which are biphasic, sustained their E peak but lost their distinct M peak, and the M-phased PAL group was now active in the evening. We next turned to imaging from DA neurons as we selectively activated different pacemaker groups that expressed ATP-gated P2X2 receptors with ATP application (46). Selective PDF neuron activation using *pdf-GAL4* excited all four DA neuron clusters that had circadian Ca^2+^ activity rhythms (PAL, PPL1, PPM1/2, and PPM3; Figure 7A-C). In contrast, the PAM cluster, which did not produce circadian Ca^2+^ activity rhythms, was reproducibly inhibited by PDF neuron activation. When we selectively activated E cells using the same method, only the DA neuron clusters that showed evening activity peaks, PPL1 and PPM3, were excited by E-cell activation; PAL and PAM were inhibited by E-cell activation (Figure 7D-F). Together these results indicated broad sensitivity of DA clusters to M cell activation and somewhat less to E cell activation. In addition, it suggested that the daily activity pattern of PAL neurons might be shaped by a combination of excitatory inputs from M cells in the morning and inhibitory inputs from E cells in the evening. Thus, many of the circadian pacemaker groups have neuronal connectivity patterns that could support them regulating the phases of activity in downstream non-pacemaker DA neurons.

**Figure 7.**
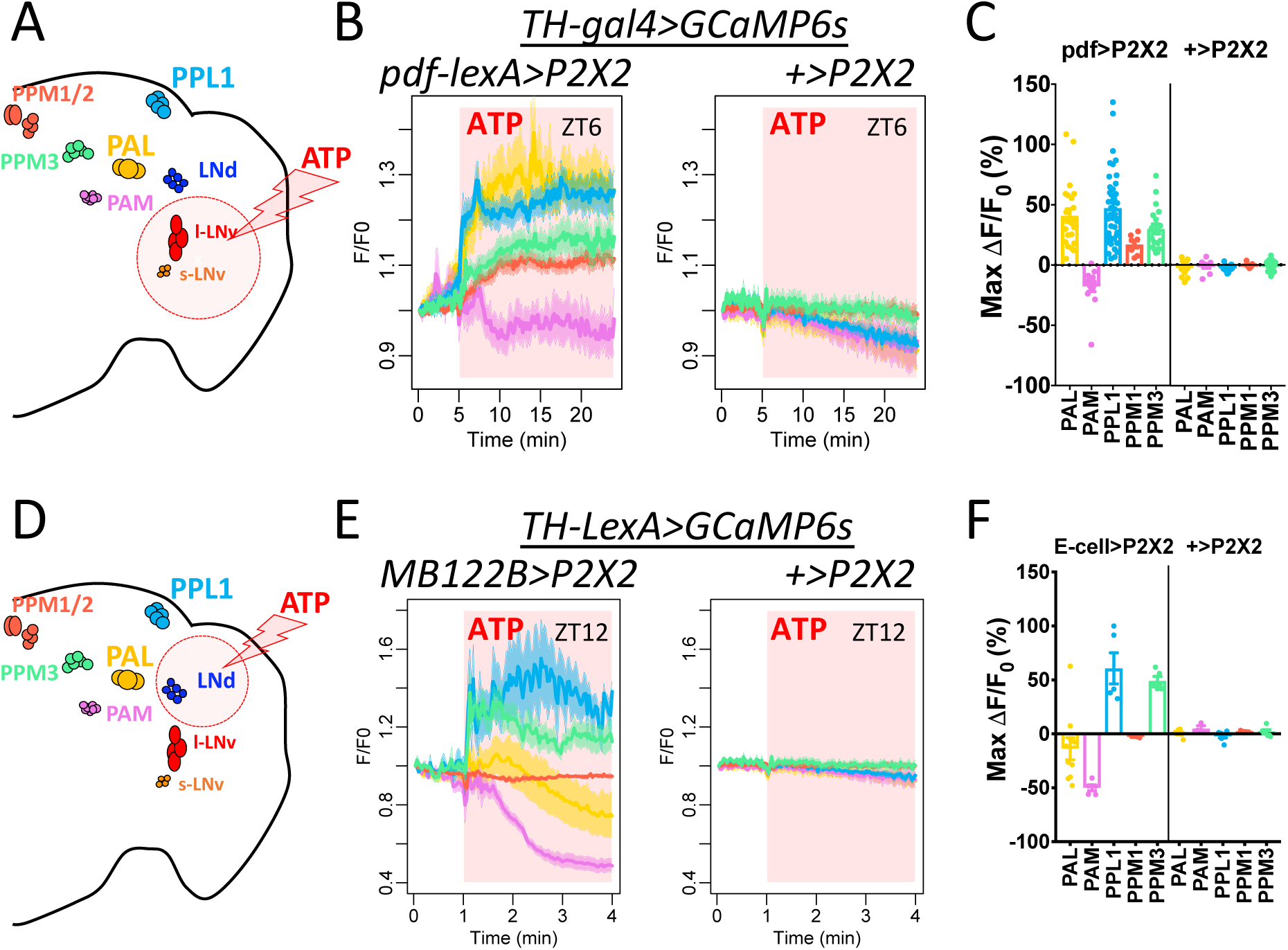
Functional connections from M and E cells DA neuron clusters. (**A**) Illustration of pharmacological activation of PDF-positive neurons (sLNv and lLNv). (**B**) Average traces of DA neuron clusters responding to ATP application in flies with P2X2 expressing in PDF neurons (left, n = 7 flies) and in control flies without P2X2 expression (right, n = 4 flies). Red area indicates duration of ATP application. (**C**) Maximum Ca^2+^ signal changes after ATP application in individual cells in (B). (**D**) Illustration of pharmacological activation of E cells (the 5^th^ sLNv and 3 LNd). (**E**) Average traces of DA neuron clusters responding to ATP application in flies with P2X2 expressing in E cells (left, n = 3 flies) and in control flies without P2X2 expression (right, n = 3 flies). (**F**) Maximum Ca^2+^ signal changes after ATP application in individual cells in (E).

## Discussion

We performed a directed search among dopaminergic (DA) and peptidergic neuroendocrine (PNE) neurons across the whole fly brain for cells that display circadian neural activity rhythms. We reasoned that subsets of these groups may exhibit circadian timing patterns, as some DA neurons relate to sleep-wake regulation (20), and because in mammals the neuroendocrine system is heavily reliant on circadian regulation (47, 48, 49). In *Drosophila*, these two different neuronal complements show diverse daily activity patterns, with different PNE and DA neural centers exhibiting activity peaks at different times of day. PPM3 DA neurons display daily bimodal rhythms and they contribute to normal locomotor activity rhythms (9). *Fru+* PAL DA neurons display a morning activity peak, which is consistent with their driving a morning-biased mating rhythm (4, 5). PPL1-dFSB DA neurons displayed an evening activity peak, which is consistent with their promotion of arousal around dusk. In the *Pars Intercerebralis* (PI), insulin-producing cells (IPCs) had activity peaks in the morning, consistent with their involvement in feeding rhythms (28, 31). Other PI PNE cells displayed daily activity rhythms that peaked around Mid-Day, and which likely underlie rhythms of hormone secretion for multiple neuroendocrine systems. Daily neural activity rhythms of these output circuits were dependent on the molecular clock and driven by activity derived in the circadian pacemaker circuit. Based on these findings, we hypothesize that multiple, sequential neuronal outputs from the polyphasic circadian pacemaker circuit are used to assign diverse phases to different physiological processes and behaviors as illustrated in Figure 8.

**Figure 8.**
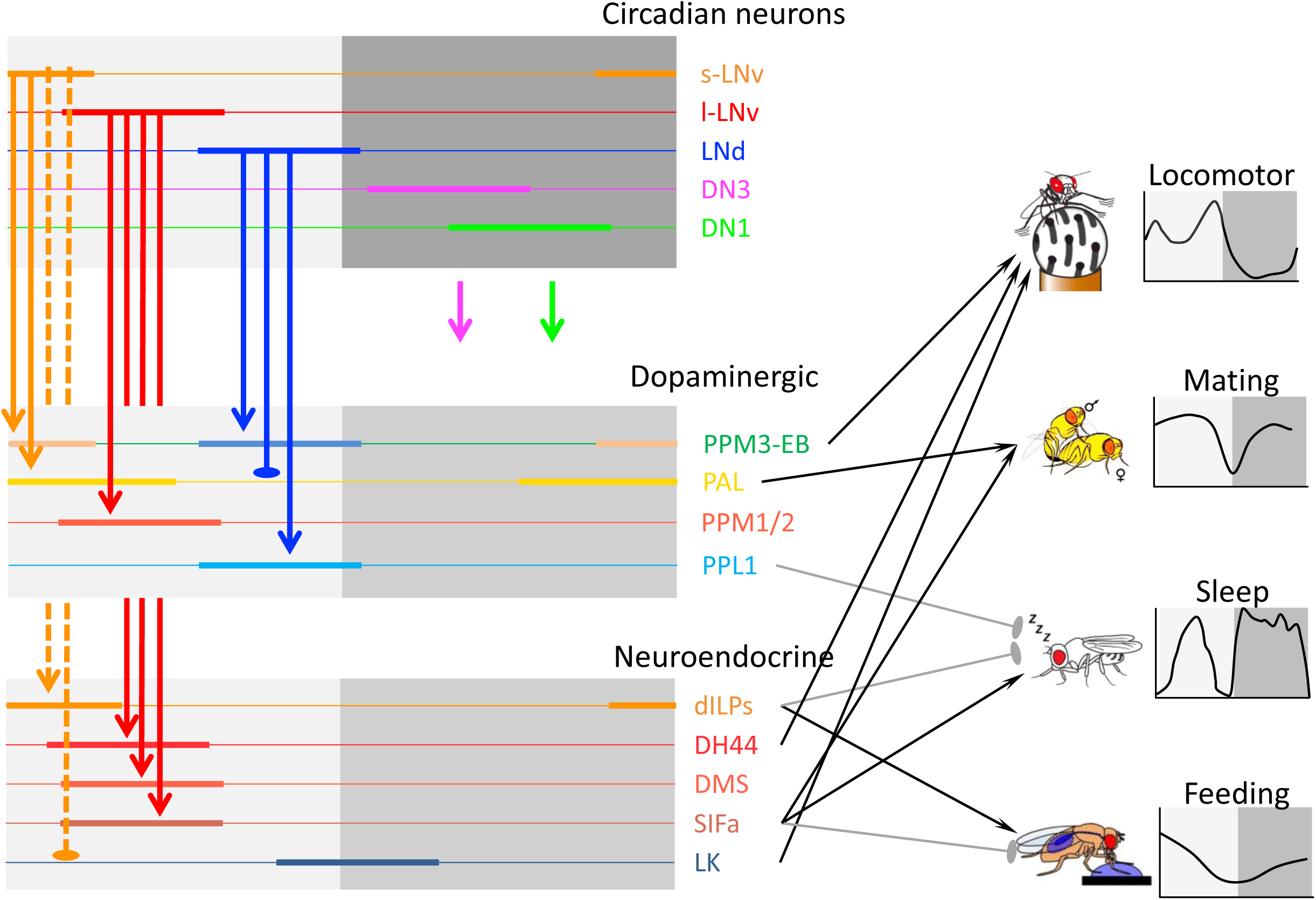
A model of the polyphasic circadian output pathways in *Drosophila*. Groups of circadian neurons peaking at different times of day sent outputs to different downstream circuits to generate diverse phases. Circadian pacemaker M cells (orange arrows) and E cells (blue arrows) independently activate PPM3 neurons around dawn and dusk, which help drive 3locomotor activity rhythms (9). Fru+ PAL neurons might be activated by M cells and inhibited by E cells and drive mating behavior (the daily mating pattern was redrawn from 4). E cells activate PPL1-dFSB neurons to suppress sleep around dusk (the daily sleep pattern was redrawn from 17). M cells activate insulin-like peptides(dILPs) producing PI neurons, which then promote (black arrow) feeding and suppress (gray arrow) sleep around dawn (the daily feeding pattern was redrawn from 3). M cells might inhibit Leucokinin (LK) neurons, which regulate locomotor rhythms and metabolisms incorporated with daily feeding rhythms. l-LNv (red arrows) controls the midday phase of dILP-negative PI neurons and PPM1 DA neurons.

We found that the spontaneous activity patterns of three distinct groups of DA neurons: PAL, PPL1, and PPM1/2 are under circadian control, similar to that displayed by the DA-PPM3 group (9). Previous studies have described synaptic connections between DA neurons and circadian neurons (50): they suggested that DA regulates circadian neuron activity. Our findings, as well as other recent studies (51, 52) argue that circadian pacemakers also regulate DA neuron activity. DA neurons responded to circadian neuron activation (Figure 7) and showed circadian neural activity rhythms (Figure 1B). DA neural activity rhythms required functional clock gene oscillations (Figure 1C) and normal circadian pacemaker neurotransmission (Figure 1D). Lastly, different phases of DA neural activity rhythms were dictated by phases of different circadian neuron groups (Figure 4-6).

In addition to rhythmicity in locomotor activity (9), circadian rhythms in DA neurons might be involved in organizing the daily sleep/wake pattern. Besides the nighttime sleep, *Drosophila* exhibit a mid-day siesta (daytime sleep) in between the morning and evening activity periods. DA promotes wakefulness and arousal (20), in part through a pair of PPL1-dFSB neurons inhibiting the sleep-promoting dFSB neurons (17, 25). We found that PPL1-dFSB neurons were active around dusk (Figure 2C), which might be driven by the output of E cells (Figure 4C). The activity pattern suggested PPL1-dFSB may selectively regulate afternoon wakefulness around the time of the evening activity peak. We speculate that the morning wakefulness might be controlled by a separate circuit, such as through neuropeptide DH31 released from DN1 neurons (53) and IPCs (Figure 3D; 28). In contrast, the mid-day siesta might be promoted by SIFa neurons in PI (Figure 3G; 29). Similarly, a recent study by Sun *et al.,* (10 2022) showed that outputs of DN1 and DN3 subsets during the night together promote nighttime sleep through mushroom body circuitry, via a group of specific cholinergic neurons.

Regarding reproductive behavior, circadian mating rhythms peak in the morning and may be driven in part by males (4, 5). Zhang *et al.* (26) showed that a different pair of DA neurons, the *fruitless*-positive (*Fru+*) PAL, encode internal mating drive in male flies by regulating P1 neurons. We found that *Fru+* PAL neurons were spontaneously active in the morning (Figure 2D) a phase consistent with the expectation that spontaneous *Fru+* PAL activity promotes mating drive in the morning. It was striking that the morning activity peak of PAL neurons, unlike that of the PPM3 and EB ring neurons, was not dictated by M-cell activity alone (Figure 4B). Instead, E-cell activity had a stronger effect in determining the phase of PAL activity peak (Figure 4C), consistent with the previous observation that impairing clocks in E cells (LNd) and in DN1 caused more severe mating rhythm deficits (54). Interestingly, E cells were in antiphase to PAL, suggesting that unlike their activation of PPL1 and PPM3 neurons, E cells might regulate the activity pattern of PAL neurons, at least in part, through inhibition (Figure 7E). A third DA neuron group PPM1/2 peaked at Mid-Day, and its phase was aligned with and sensitive to the phase of l-LNv activity (Figure 5-6). However, no specific functions for PPM1/2 have as yet been described.

Neuropeptides released by PNE cells regulate multiple aspects of *Drosophila* physiological states and behaviors (27). We found several groups of PNE cells that exhibit circadian neural activity rhythms, including those expressing dILP2, SIFa, DMS, and DH44 in the PI, and LK neurons in lateral horns (Figure 3). dILP2 neurons (a.k.a. insulin-producing cells, IPC), which promote feeding and suppress sleep (28, 31), peaked in the morning and may be controlled by M cells (Figure 3D; 55). The other PI neurons peaked around Mid-Day, including the SIFa, DMS, and DH44 neurons. SIFa neurons can promote sleep (29) and mating (56), and also suppress feeding (32). DH44 neurons together with a pair of LK neurons regulate locomotor activity rhythms (30, 38) LK neurons are also involved in metabolism and regulate behavior associated with daily feeding rhythms (35, 57). We found that DH44 neuron activity peaked around Mid-Day, whereas that of LK neurons peaked in early evening (Figure 3): these data are consistent with the activity patterns of these two groups of peptidergic neurons when measured previously in acutely dissected brains (38). Together, dILP2, SIFa, and LK neurons, with different activation phases and effects, help shape the daily feeding pattern (Figure 8; 3). However, the activity patterns of DH44 and LK neurons were different from the profile of locomotor activity. Further studies are required to determine how the DH44 and LK neuronal activity patterns specifically contribute to the daily bimodal pattern of locomotor rhythms.

More generally, these studies prompt consideration of how polyphasic circadian timing information is normally transmitted from clock-expressing pacemakers to non-clock-expressing “downstream neurons”. In mammals, numerous hormones are released in circadian patterns and at different times of day. For example, melatonin is uniformly released in the night, while glucocorticoids are normally released in anticipation of waking, a phase-point that varies widely among different species. Moreover, circadian regulation over daily hormone release depends on direct connectivity with the neurons of the suprachiasmatic nucleus (SCN) (59, 60). Jones *et al.* (61) recently studied circadian corticosterone production and showed that VIP-secreting neurons of the SCN delay corticotropin-releasing hormone (CRH) release by inhibiting CRH neurons of the paraventricular nucleus. The inhibition is two-fold: VIP neuron activation entrains the Period-based molecular clock intrinsic to the CRH neurons. In addition, VIP neurons acutely suppress CRH neuron activity by depressing basal Ca^2+^ levels. The latter is a phenomenon very similar to the effect of neuropeptide PDF in *Drosophila*: PDF suppresses neuronal activity in the LNd Evening pacemakers by depressing their basal Ca^2+^ levels for many hours (8).

We term the *Drosophila* pacemaker system “polyphasic” because its constituent neural groups produce at least five distinct and stereotyped phases of neuronal activity across the solar day: the M, MD, E, N (Night)1 and N2 phases (7). Different subsets of DA and PNE neurons exhibit similar polyphasic activity patterns with different subsets aligning unambiguously with the different phases of the pacemaker network. For example (i) M phase activity is displayed by DA-PAL and PNE-dILP2, (ii) E phase activity is displayed by the DA-PPL1 and PNE-LK, (iii) both M and the E phase activity is displayed by the DA-PPM3, or (iv) the MD phase is displayed by the DA-PPM1/2, PNE-SIFa, PNE-DMS, and PNE-DH44. The simplest hypothesis would suggest a one-to-one relationship between the driver for a particular circadian phase and the followers for that phase point. To some extent, there is support for that possibility: the M and E pacemakers independently regulate the morning and evening phases of activity in the biphasic DA-PPM3 and EB-RNs (9). However, in other cases, phasic control may be more complex: Here we found that the DA-PAL is normally active in the morning and aligned with M (s-LNv) pacemakers. However, advancing the phase of either the M or the E pacemakers advanced the PAL phase (Figure 4), suggesting the PAL morning phase is normally the product of at least two different sources of pacemaker input. We found complexity also in the regulation of MD-active downstream neurons. The MD phase point is represented by the activity of the l-LNv and its ability to control the phase of neurons normally active at MD was shown by experimental manipulation (Figure 5D) and importantly also by tracking normal developmental progression (Figure 6). The l-LNv are themselves PNE neurons that secrete the neuropeptide PDF. There is at present no strong evidence to support the possibility of additional l-LNv transmitters, suggesting PDF is the basis by which the MD phase is relayed from the pacemaker system to downstream centers in the instances we documented MD phase alterations. However, with loss of PDF signaling (as measured in a *pdfr* gene mutation in mature *Drosophila*), the MD phase remains intact (Figure 1D). Hence the cellular-molecular basis that defines the MD phase in the mature adult remains enigmatic, both within the pacemaker circuit (7) and outside it. Both are insensitive to loss of function for PDF signaling, yet both respond with multi-hr phase delays to greater PDFR expression by l-LNv.

Irrespective of its basis, our results clearly show that the MD (mid-day) timepoint is a third *bona fide* phase marker produced by the circadian pacemaker circuit. This finding extends the definition of functional neuronal oscillators in *Drosophila* beyond the two canonical Morning and Evening ones (e.g., 7, 9; 42, 62, 63, 64). In summary, we found multiple neural pathways relating the circadian pacemaker system with daily rhythms of behaviors. Different groups of circadian neurons, acting alone and/or in concert, impose diverse neural activity rhythms onto different groups of downstream DA and PNE neurons. These downstream neurons then separately or synergistically regulate the daily rhythms in locomotor activity, sleep/wake, feeding, and mating behaviors. Notably, several groups of downstream neurons have been suggested to be involved in the interaction between different rhythmic behaviors - sleep and mating (65, 66), and sleep and feeding (35, 67). Our findings suggest parallel and over-lapping control from circadian neurons to downstream functional circuits which may be a substrate to regulate such interactions. Future studies will help to define the precise nature of the cellular and molecular signals by which the polyphasic circadian timing system is translated across a wide array of physiological outputs.

## Acknowledgments

We thank Holy and Taghert laboratories and the Washington University Center for Cellular Imaging (WUCCI) for advice and technical support; Mark Wu, Margaret Ho, and Tingting Xie for guidance in selecting driver lines for dopaminergic neurons; Orie Shafer, Francois Rouyer, Larry Zipursky, Michael Rosbash, and Bloomington Stock Center provided fly stocks and reagents. The work was supported by the Washington University McDonnell Center for Cellular and Molecular Neurobiology and by NIH grants R01 NS068409 and R01 DP1 DA035081 (T.E.H.), R01 NS099332 and R01 GM127508 (P.H.T.), and R24 NS086741 (T.E.H. and P.H.T.).

## Author Contributions

X.L., T.H.E., and P.H.T. conceived the experiments; X.L. performed and analyzed all experiments; X.L., P.H.T. and T.H.E. wrote the manuscript.

## Supplemental figure legends

**Figure S1.**
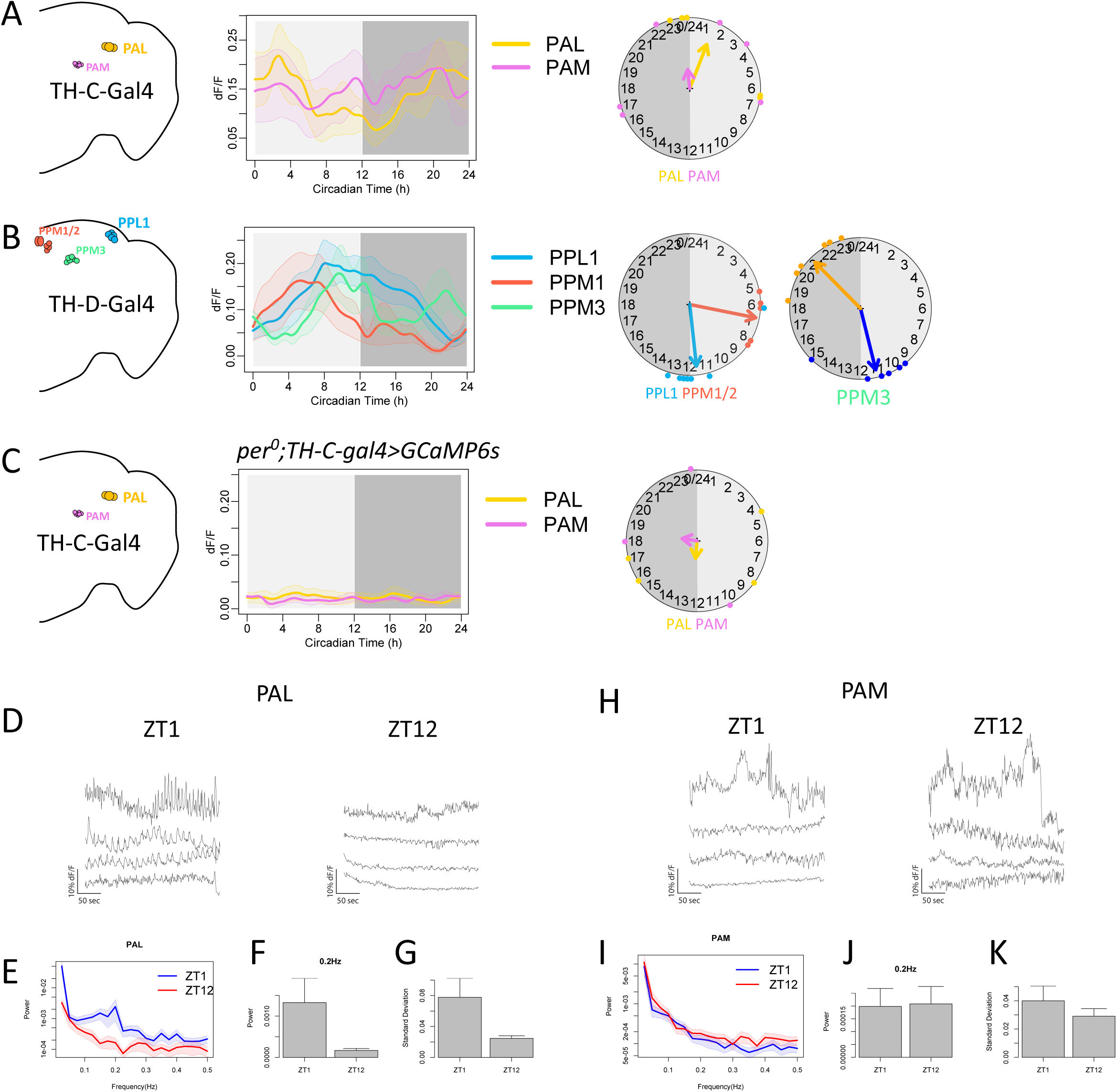
Diverse daily Ca^2+^ activity patterns of DA neuron clusters. (**A**) Left, map of the two DA neuron clusters: PAL and PAM, accessible via in vivo imaging labelled by *TH-C-Gal4*. Right, daily Ca^2+^ activity patterns of these two DA neuron clusters under DD (n = 6 flies). (**B**) Left, map of the three DA neuron clusters: PPL1, PPM1/2, and PPM3, labelled by *TH-D- Gal4*. Right, daily Ca^2+^ activity patterns of these three DA neuron clusters under DD (n = 6 flies). (**C**) Arrhythmic Ca^2+^ activity patterns of two DA neuron clusters labelled by *TH-C-Gal4* under DD in *per^01^* mutants (n = 5 flies). (**D-K**) Fast Ca^2+^ activity of two DA clusters: PAL (D-G) and PAM (H-K), measured by 1Hz imaging at two zeitgeber time points: ZT1 (n = 7 flies) and ZT12 (n = 5 flies). (**E and I**) Power spectrums (fast Fourier transform) of PAL and PAM activity at two ZTs. (**F**) For PAL, power at 0.2Hz is stronger in the morning than in the evening. (**J**) For PAM, power at 0.2Hz is not different between morning and evening. (**G**) For PAL, overall standard deviation is higher in the morning than in the evening. (**K**) For PAM, overall standard deviation is not different between morning and evening.

**Figure S2.**
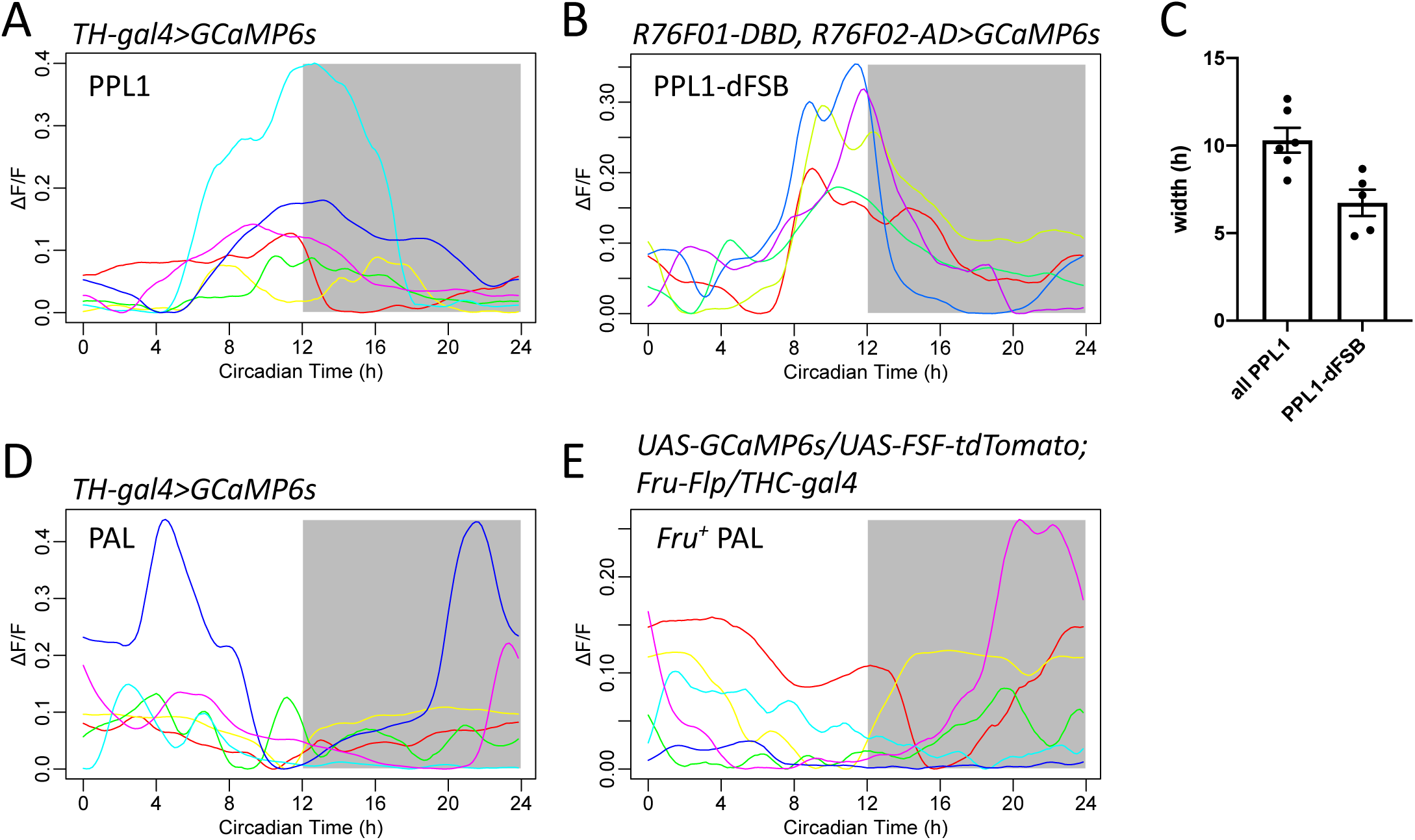
Daily Ca^2+^ activity traces of subgroups of DA neuron clusters. (**A**) Activity traces of PPL1 neurons from Figure 1B. Individual flies are plotted in different colors. (**B**) Individual activity traces of PPL1-dFSB subgroup from Figure 2C. (**C**) Comparison of peak widths (the full width at half maximum) for Ca^2+^ transients (t test, p = 0.0074). (**D**) Individual activity traces of PAL neurons from Figure 1B. (**E**) Individual activity traces of *Fru+* PAL subgroup from Figure 2D.

**Figure S3.**
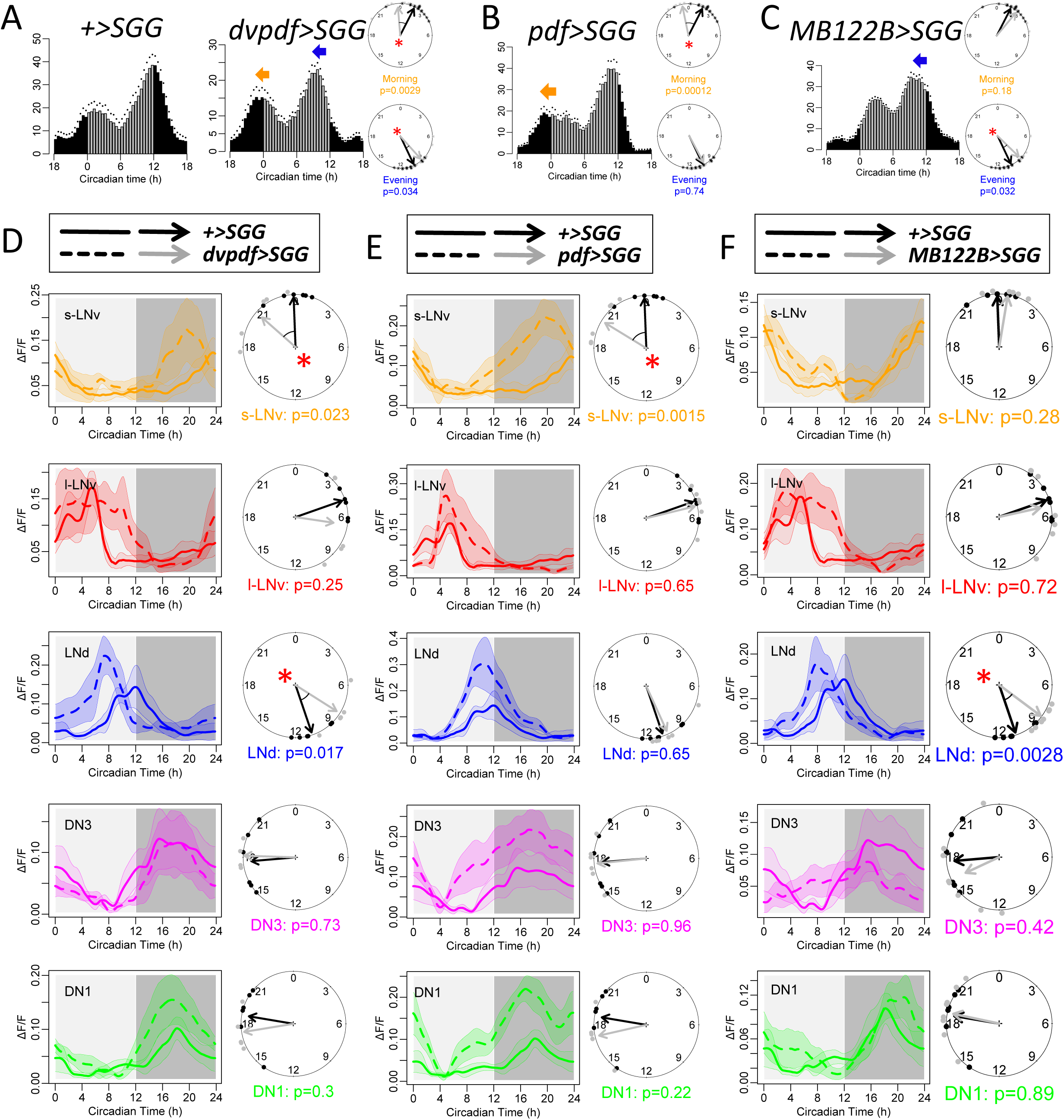
Daily activity phases of DA neurons are dictated by different groups of circadian neurons. (**A**) Average locomotor activity in DD1 of wild type flies (WT, n = 16) and of flies expressing *SGG* in PDF neurons and E cells using *dvpdf-GAL4* (*dvpdf>SGG*, n = 16 flies). Phase comparisons of morning and evening activity between WT and *dvpdf>SGG*. (**B**) Average locomotor activity in DD1 of flies expressing *SGG* in PDF neurons using *pdf-GAL4* (*pdf>SGG*, n = 16). Note that only the morning activity phase was advanced (* P < 0.05, Watson-Williams test). (re-plotted from 9) (**C**) Average locomotor activity in DD1 of flies expressing *SGG* in E cells using split-GAL4s *MB122B* (*MB122B>SGG*, n = 16). Note that only the evening activity phase was advanced (* P < 0.05, Watson-Williams test). (re-plotted from 9) (**D**) Daily Ca^2+^ activity patterns and phase comparison of circadian neurons between WT flies under DD (solid lines, n = 4 flies) and *dvpdf>SGG* flies under DD (n = 6 flies). The daily peak phases of s-LNv and LNd were advanced in *dvpdf>SGG* flies (* P < 0.05, Watson-Williams test). (**E**) Daily Ca^2+^ activity patterns and phase comparison of circadian neurons between WT flies under DD (solid lines, n = 4 flies) and *pdf>SGG* flies under DD (dashed lines, n = 5 flies). The daily peak phases of s-LNv were significantly advanced in *pdf>SGG*. (re-plotted from 9) (**F**) Daily Ca^2+^ activity patterns and phase comparison of circadian neurons between WT flies under DD (solid lines, n = 4 flies) and *MB122B>SGG* flies under DD (n = 6 flies). The daily peak phases of LNd were advanced in *MB122B>SGG* flies (* P < 0.05, Watson-Williams test). (Data re-plotted from 9)

## STAR Methods

### KEY RESOURCES TABLE

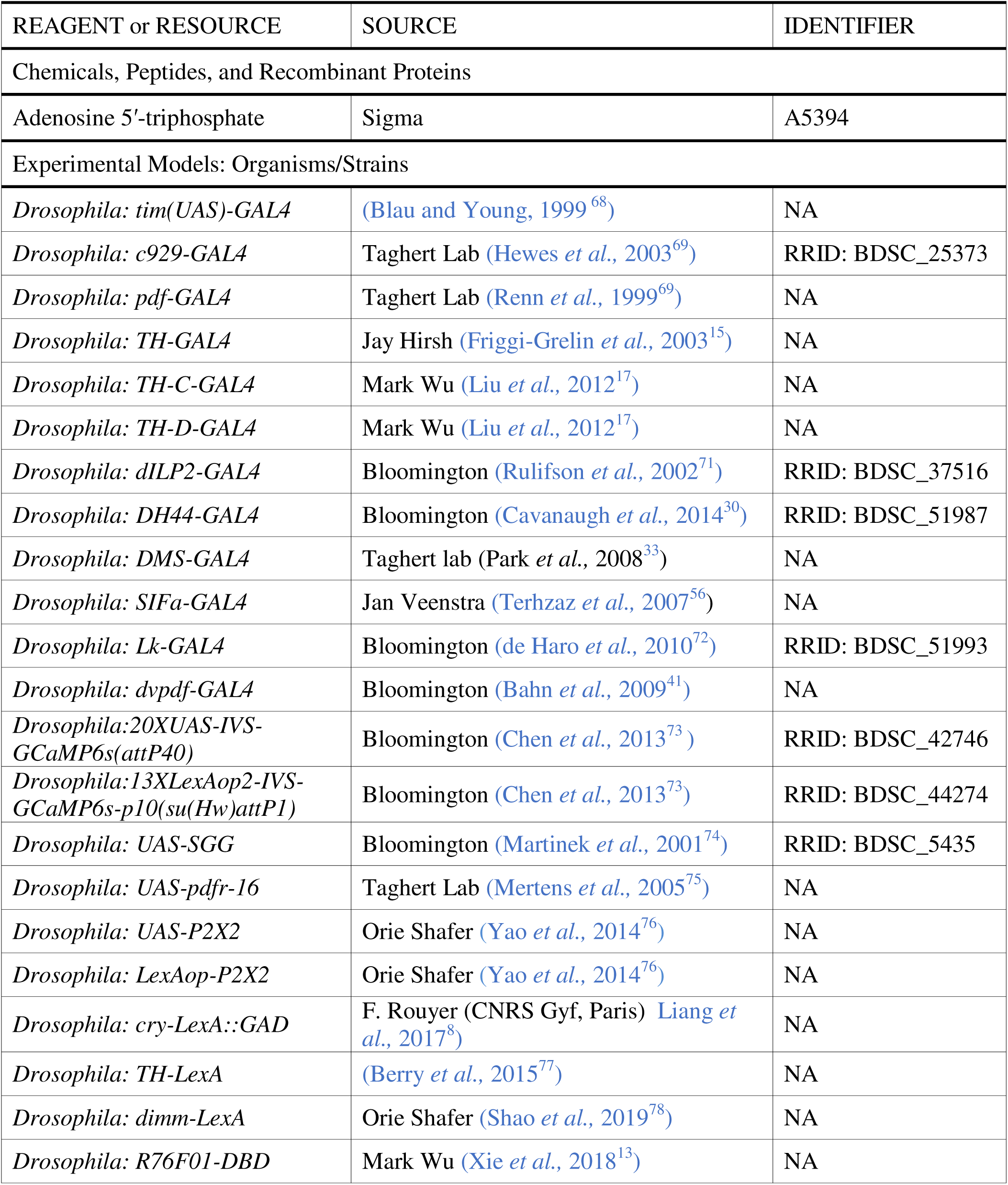
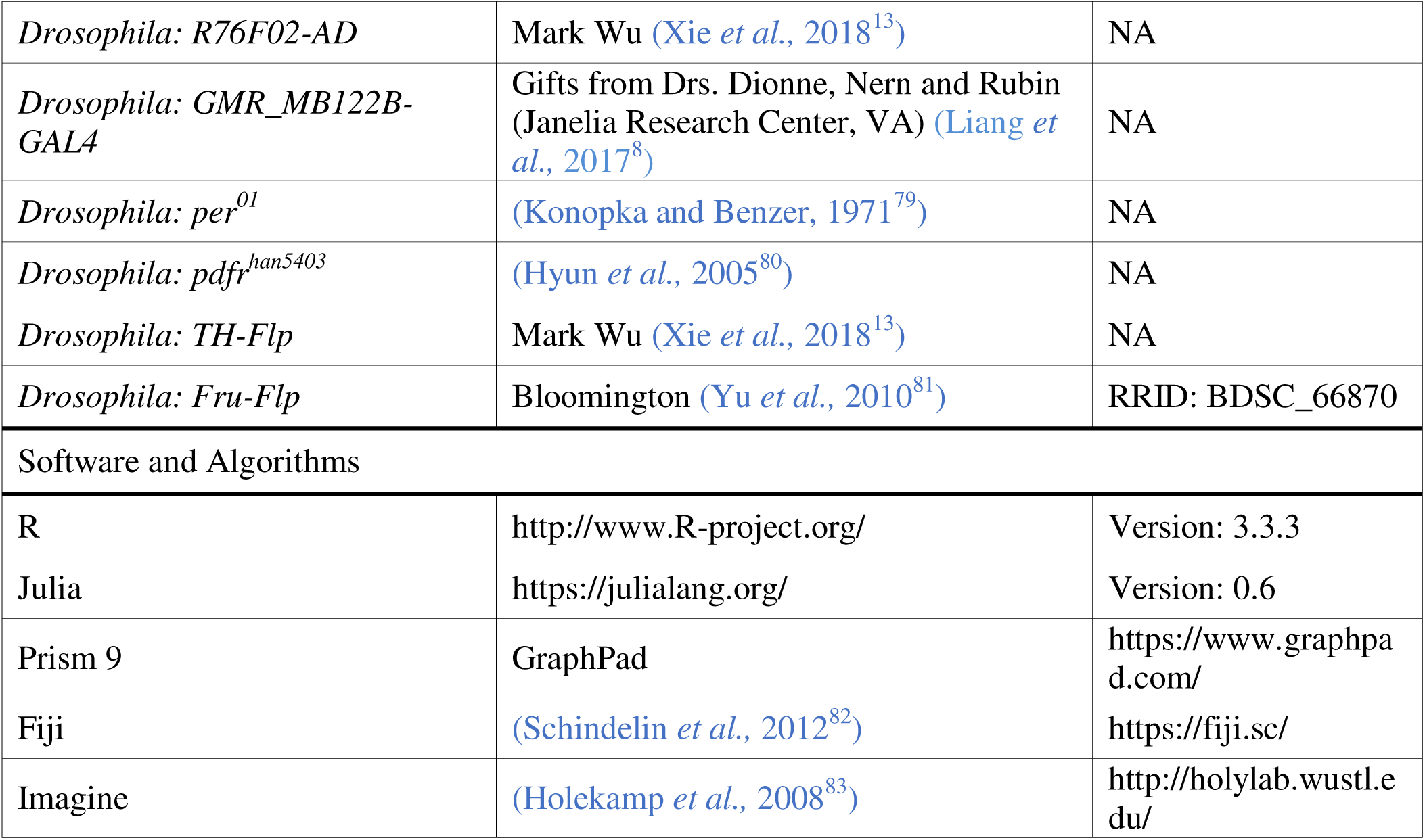

### CONTACT FOR REAGENT AND RESOURCE SHARING

Further information and requests for reagents may be directed to and will be fulfilled by the Lead Contact, Paul H. Taghert.

### EXPERIMENTAL MODEL AND SUBJECT DETAILS

#### Fly stocks

Flies were reared on standard cornmeal/agar food at room temperature. Before imaging experiments, flies were entrained under 12 h light: 12 h dark (LD) cycles at 25°C for at least 3 days. All experiments used male flies older than three days after eclosion except for Figure 6, in which flies within one day after eclosion were used.

The following fly lines have been described previously: *tim(UAS)-GAL4* (68), *TH-GAL4* (15), *TH-C-GAL4* and *TH-D-GAL4* (17), *dILP2-GAL4* (71), *DH44-GAL4* (30), *DMS-GAL4* (33), *SIFa-GAL4* (56), *Lk-GAL4* (72), *c929-GAL4* (69), *dvpdf-GAL4* (41); split-GAL4 lines: *R76F01-DBD* and *R76F02-AD* (13); *TH-LexA* (77), *dimm-LexA* (78), *cry-LexA* (8), *pdf-LexA* (22); *Fru-Flp* (81); *UAS-SGG* (42)*, UAS-pdfr* (75)*, UAS-P2X2* and *LexAop-P2X2* (76), *UAS-GCaMP6s* and *LexAop-GCaMP6s* (73); *per^01^* (79) and *pdfr^han5403^* (80).

The *cry-LexA* line was a gift from Dr. F Rouyer (CNRS Gyf, Paris). The *UAS-(FRT.stop)- tdTomato* line was a gift from Dr. S. Lawrence Zipursky (UCLA). The *dimm-LexA* line was a gift from Jim Truman (Janelia Research Center, VA).

### METHOD DETAILS

#### *In vivo* fly preparations

The surgical procedures were as previously described (7; 9). The flies were mounted by inserting the neck into a narrow cut in a piece of aluminum foil. Thus, the foil separated the head from the body and permitted the immersion of exposed brain by saline, while leaving the body in an oxygen-normal environment. During brain-exposing surgery and in vivo imaging, the head was immersed in saline, while the body remained in an air-filled enclosure. To access circadian neurons, one antenna, a portion of the dorso-anterior head capsule, and a small part of one compound eye were removed. To access dopaminergic (DA) neurons and Pars Intercerebralis (PI), a portion of the dorsal head capsule and the ocelli were removed, while the compound eyes and antennae remained intact. The orientation of fly was then tilted for a more optimal view of DA neurons in the posterior brain.

#### *In vivo* calcium imaging

For long-term (24-hr) *in vivo* imaging, a custom horizontal-scanning Objective Coupled Planar Illumination (OCPI) microscopes (83) were used, as described in Liang *et al.* (7; 8). Briefly, the ∼5 µm thick light sheet was scanned across the fly brain through a small cranial window every 10 min with a step size of 10 microns to acquire 20 to 40 separate images. Exposure time of each image was less than 0.1 s. For short-term high-frequency imaging, a custom high-speed OCPI-2 microscope (84) was used, acquiring volumetric images with 0.1-1Hz, as described in Liang *et al.* (9; 18). All imaging was performed under constant darkness and fresh HL3 saline (5 mM KCl, 1.5 mM CaCl2, 70 mM NaCl, 20 mM MgCl2, 10 mM NaHCO3, 5 mM trehalose, 115 mM sucrose, and 5 mM HEPES; pH 7.1) was perfused continuously (0.1-0.2 mL/min). For pharmacological tests, after 5-min baseline recordings, 10 mM ATP solution (pH was adjusted to 7) was manually added into a 9 ml static HL3 bath over a ∼2 s period.

#### Data reporting

No statistical methods were used to predetermine sample sizes. The selection of flies from vials for imaging and behavioral tests were randomized. The investigators were not blinded to fly genotypes.

### QUANTIFICATION AND STATISTICAL ANALYSIS

#### Imaging data analysis

Calcium imaging data was analyzed as described previously (7; 9; 18). Images were acquired by custom software (Imagine – 84) and processed in Julia 0.6, including non-rigid registration, alignment and maximal projection along z-axis. Then ImageJ-based Fiji (82) was used for rigid registration and to manually select regions of interest (ROIs) over individual cells or groups of cells. Average intensities of ROIs were measured through the time course and divided by the average of the whole image to subtract background noise.

For spontaneous calcium transients, each time trace was calculated as dF/F=(F-F_min_)/F_mean_. For 24-hr time traces, traces of certain cell type ROIs were firstly aligned based on Zeitgeber Time and then averaged across different flies. The phase relationship between traces was estimated by cross-correlation analysis. The 24-hr-clock circular plot of phases reflected both mean peak time and phase relationship of the same cell-group traces from different flies. For neurons with daily bimodal patterns (PPM3 DA neurons), each trace was split into two parts: ZT18-ZT6 (morning) and ZT6-ZT18 (evening) to estimate the morning and evening peak phases respectively. For pharmacological calcium responses, each time trance was normalized by the initial intensity (F/F_0_). The maximum change was calculated by the maximum difference of normalized intensities between baseline and following drug application. Trace analysis and statistics were performed using R 3.3.3 and Prism 7 (GraphPad, San Diego CA).

